# Capsaicin alters human Na_V_1.5 mechanosensitivity

**DOI:** 10.1101/2021.07.13.452086

**Authors:** Luke M. Cowan, Peter R. Strege, Radda Rusinova, Olaf S. Andersen, Arthur Beyder, Gianrico Farrugia

**Affiliations:** Enteric Neuroscience Program (ENSP), Division of Gastroenterology and Hepatology, Mayo Clinic, Rochester, MN; Department of Physiology and Biomedical Engineering, Mayo Clinic, Rochester, MN; Department of Physiology and Biophysics, Weill Cornell Medical College, New York, NY

**Keywords:** amphipathic, arrhythmia, capsaicin, electrophysiology, functional gastrointestinal disorder, ion channel, irritable bowel syndrome, mechanosensitivity, mechanotransduction, voltage-gated sodium channel type 5

## Abstract

*SCN5A*-encoded Na_V_1.5 is a voltage-gated Na^+^ channel expressed in cardiac myocytes and human gastrointestinal (GI) smooth muscle cells (SMCs). Na_V_1.5 contributes to electrical excitability in the heart and slow waves in the gut. Na_V_1.5 is also mechanosensitive, and mechanical force modulates several modes of Na_V_1.5’s voltage-dependent function. Na_V_1.5 mutations in patients with cardiac arrhythmias and gastrointestinal diseases lead to abnormal mechano- and voltage-sensitivity. Membrane permeable amphipathic drugs that target Na_V_1.5 in the heart and GI tract alter Na_V_1.5 mechanosensitivity (MS), suggesting that amphipaths may be a viable therapeutic option for modulating Na_V_1.5 function. We therefore searched for membrane-permeable amphipathic agents that would modulate Na_V_1.5 MS with minimal effect on Na_V_1.5 voltage-gating intact to more selectively target mechanosensitivity. We used two methods to assess Na_V_1.5 MS: (1) membrane suction in cell-attached macroscopic patches and (2) fluid shear stress on whole cells. We tested the effect of capsaicin on Na_V_1.5 MS by examining macropatch and whole-cell Na^+^ current parameters with and without force. The pressure- and shear-mediated peak current increase and acceleration were effectively abolished by capsaicin. Capsaicin abolished the mechanosensitive shifts in the voltage-dependence of activation (shear) and inactivation (pressure and shear). Exploring the recovery from inactivation and use-dependent entry into inactivation, we found divergent stimulus-dependent effects that could potentiate or mitigate the effect of capsaicin, suggesting that mechanical stimuli may differentially modulate Na_V_1.5 MS. We conclude that selective modulation of MS makes capsaicin is a novel modulator of Na_V_1.5 MS and a promising therapeutic candidate.

## INTRODUCTION

The S*CN5A*-encoded voltage-gated sodium channel Na_V_1.5 is mechanosensitive; mechanical force modulates Na_V_1.5’s voltage-dependent function. This property is particularly relevant given that Na_V_1.5 is expressed in mechanically active tissues like the heart and gastrointestinal tract, where it contributes to electrical excitability in cardiac myocytes (CMs) and slow waves in smooth muscle cells (SMCs), respectively.^1^ At the cellular level, mechanosensitive channels like Na_V_1.5 detect mechanical stimuli through lipid bilayer tension and/or cytoskeletal deformation.^2,3^ In patch-clamp studies, mechanical stimuli in the form of membrane stretch and fluid shear stress modulate Na_V_1.5’s voltage-dependent function by increasing whole-cell conductance, shifting voltage-dependence to hyperpolarized potentials, and accelerating its activation and inactivation kinetics. Therefore, in the context of cardiac and intestinal smooth muscle tissues, Na_V_1.5 mechanosensitivity (MS) has important implications.^1^

Channel-related disorders in common cardiac and gastrointestinal diseases, channelopathies^*4, 5*^ are strongly linked to *SCN5A* mutations.^*6, 7*^ Many *SCN5A* mutations lead to voltage-gating abnormalities, some associated with abnormal responses to mechanical stimuli.^*7-10*^ Impaired stretch modulation in Na_V_1.5, for example, occurs in some mutations that also cause LQT3-type cardiac arrhythmias,^*8, 11*^ and some mutations that lead to altered Na_V_1.5 MS are found in patients with IBS.^*10, 12*^

Because of their involvement in many diseases, ion channels are prime targets for pharmacological treatment.^13^ Drugs for cardiac diseases may modulate the behavior of sodium channels like Na_V_1.5 and influence mechanosensitivity.^13-15^ Yet, how membrane-permeable amphipathic drugs alter the mechanosensitivity of voltage-gated channels like Na_V_1.5, or whether they do so by a mechanism separate from Na^+^ current inhibition, remain critical unanswered questions.^13-15^ For example, the membrane-permeable, amphipathic local anesthetic lidocaine inhibits peak current while inhibiting Na_V_1.5 mechanosensitivity at lower concentrations.^*13, 14*^ Suggesting separate mechanisms for current inhibition and altered mechanosensation by amphipaths, the anesthetic binding site mutation F1760A^13^ eliminates the voltage-dependent inhibition by lidocaine without altering lidocaine’s effect on mechanosensitivity. The membrane-impermeant lidocaine analog, QX-314, in contrast had no effect on mechanosensitivity.

We therefore searched for membrane-permeable amphipathic agents with minimal effects on Na_V_1.5 voltage-gating for their impact on Na_V_1.5 mechanosensitivity. Among the candidates, capsaicin shows promise, and we characterized its ability to selectively modulate Na_V_1.5 mechanosensitivity.

## METHODS

### Heterologous expression and cell culture

Wild-type *SCN5A* (Q1077del Na_V_1.5)^16^ was co-transfected with pEGFP-C1 into HEK-293 cells with Lipofectamine 3000 reagent (Thermo Fisher Scientific, Massachusetts, USA).

### Electrophysiology

#### Pipette fabrication

For whole-cell experiments, electrodes were pulled on a P-97 puller (Sutter Instruments, CA) from KG12 glass to a resistance of 2-5 MΩ. For cell-attached patch experiments, electrodes were pulled from 8250 glass (King Precision Glass, California, USA) then fire-polished to wide-bore, bullet-shaped tips with a final resistance of 1-2 MΩ. Electrodes were coated with R6101 elastomer (Dow Corning, MI) then cured by a heat gun to reduce capacitive transients.

#### Data acquisition

Whole-cell and cell-attached patch data from HEK-293 cells were recorded at 20 kHz with an Axopatch 200B patch-clamp amplifier, Digidata 1550, and pClamp11 software (Molecular Devices, CA).

#### Cell-attached patch

*Solutions*: The pipette solution contained (in mM): 150 Na^+^, 160 Cl^-^, 5 K^+^, 2.5 Ca2^+^, 10 HEPES, and 5.5 glucose with an osmolality of 305 mmol/kg and pH of 7.35. GdCl_3_ (10 µM) was included in the pipette solution to inhibit endogenous stretch-activated channels. The bath solution contained (in mM): 15 Na^+^, 140 Cs^+^, 160 Cl^−^, 2.5 Ca^2+^, 5 K^+^, 10 HEPES, and 5.5 glucose with an osmolality of 305 mmol/kg and pH of 7.35. Where applicable, capsaicin was diluted 1000-fold in bath solution from a 20 mM ethanol stock then added to the recording chamber. Seal pressures were digitally controlled and monitored by High-Speed Pressure Clamp (HSPC-2, ALA Scientific, NY). Suction ≤10 mmHg was applied to establish giga-seals. *Episodic protocol and mechanical stimulation by pressure*. Na^+^ currents in macroscopic patches were elicited by an identical pair of voltage ladders with 31-ms pressure steps up to −50 mmHg encompassing the second voltage ladder. Patches were held at +100 mV, stepped briefly for 10 ms to +190 mV to close Na_V_ channels, then stepped through a 10-step voltage ladder from +100 to 0 mV in 21-ms long, 10-mV increments with a total sweep duration (equivalent to the interpulse interval) of 280 ms. Recordings were an average of 5 runs. Capsaicin (20 µM) was added to the chamber 5 min before testing the effects of the drug. *Recovery from inactivation:* To test the effect of pressure on the recovery of Na_V_1.5 from inactivation, cells were held at 120 mV and stepped to (1) 20 mV for 30 ms, next to (2) 120 mV for a variable duration to recover, then to (2) 20 mV for 30 ms. The time between the beginning of each sweep was 5 s. The length of the recovery time in stage (2) was varied between 1 and 300 ms, with half-log unit increments. The pressure step per sweep was 400 ms regardless of recovery time. *Use-dependent inactivation:* To test the effect of pressure on the onset (use dependence) of Na_V_1.5 inactivation, cells were held at 120 mV and stepped 20 times to 20 mV, and the frequency between steps was 33.33 to 3.33 Hz. The pressure step per sweep was 30 ms.

#### Whole-cell voltage clamp

*Solutions*: The intracellular solution contained (in mM): 145 Cs^+^, 125 CH_3_SO_3-_, 35 Cl^-^, 5 Na^+^, 5 Mg^2+^, 10 HEPES, and 2 EGTA with an osmolality of 290 mmol/kg and pH of 7.0. The extracellular solution contained (in mM): 15 Na^+^, 140 Cs^+^, 160 Cl^-^, 2.5 Ca^2+^, 5 K^+^, 10 HEPES, and 5.5 glucose with an osmolality of 290 mmol/kg and pH of 7.35. *Peak current, voltage dependence of activation, and kinetics of activation and inactivation*: To measure peak Na^+^ current density, cells transfected with Na_V_1.5 were held at −120 mV then pulsed through a 2-stage, 19-step voltage ladder (1) from −110 to −30 mV in 5 mV intervals for 2.9 s each and (2) to −30 mV for 100 ms. The time from the start of each sweep to the next was 5 s. Peak currents at each voltage step were normalized to either the cell capacitance (pF) or the maximum peak inward current without shear. *Recovery from inactivation:* Recovery from inactivation was measured by holding cells at −130 mV and the pulsed through a 3-stage, 10-step protocol to (1) −30 mV for 100 ms, next to (2) −130 mV for a variable duration to recover, then to (2) −30 mV for 100 ms. The time between sweep starts was 2.5 s. The length of the recovery time in stage (2) of sweep *n* was 4*2^*n*^ ms for a total of *n* = 10 sweeps. *Use-dependent inactivation:* To measure the onset of Na_V_1.5 inactivation, cells were held at −130 mV and stepped 10 times to −40 mV, in which the frequency of steps recorded ranged between 0.3 and 50 Hz. *Mechanical stimulation by shear stress:* When testing the effect of shear stress, the extracellular (bath) solution was perfused by gravity drip (at 10 mL/min) for the duration of the voltage protocol.

### Data Analysis

The maximum peak Na^+^ current and voltage dependence of activation were determined by fitting the Na_V_1.5 current-voltage (I-V) plots with *I*_*V*_*=G*_*MAX*_*\*(V-E*_*REV*_*)/(1+e*^*(V-V1/2A)/slope*^), where *G*_*MAX*_ is the maximum Na^+^ conductance in whole cells (*I*_*MAX*_ is the maximum Na^+^ current in patches), and *V*_*1/2A*_ is the voltage of half-activation. Activation kinetics were determined by fitting currents with a two-term weighted exponential function: *I(t)=A*_*1*_*e*^*(-t/τA)*^*+ A*_*2*_*e*^*(-t/τI)*^, where *τ*_*A*_ and *τ*_*I*_ are the time constants of activation and inactivation, respectively, and *A*_*X*_ and *C* were constants. Steady-state inactivation was obtained by fitting available peak Na^+^ currents with a 3-parameter sigmoid curve: *I*_*V*_*=1/(1+e*^*((V-V1/2I)/dVI)*^), where *V*_*1/2I*_ is the half-point of steady-state inactivation (availability), and *δV*_*I*_ the slope. To determine the recovery from inactivation, peak Na^+^ currents were fit with the equation: *I/I*_*0*_*=1/(1+t/t*_*1/2*_)^*b*^, where *I/I*_*0*_ is the ratio of Na^+^ current recovered following inactivation from the control current, *b* the rate of inactivation recovery, and *t*_*1/2*_ the time where half of the Na^+^ current has recovered from inactivation. Use-dependent inactivation was estimated by fitting the peak Na^+^ currents of successive pulses with the equation: *I*_*10*_*/I*_*1*_*=I*_*f*_*e*^*b/(x+c)*^, where *I*_*10*_*/I*_*1*_ is the peak Na^+^ current in step 10 normalized to the peak of step 1, and *I*_*f*_ the maximally inactivated peak Na^+^ current at frequency *f*, and *b* or *c* is the rate or constant of use-dependent inhibition, respectively. To calculate the half-frequency of use-dependent inactivation, *I*_*f*_ was plotted *vs*. pulse frequency *f* and fit with *I*_*f*_*=(1-a)/(1+e*^*(f1/2-f)/b*^), where *a* is limit of use-dependent inhibition, *f*_*1/2*_ the half frequency of use-dependent inhibition, and *b* the slope. Data are expressed as the mean ± standard error of the mean (SEM). Significance was tested using a 2-way ANOVA with Tukey post-test, in which *P*<0.05 when comparing force to rest or capsaicin to drug-free.

## RESULTS

### Screen of amphipathic membrane-permeable drugs

We examined select amphipathic agents with high partition coefficients (Table 1) as potential modulators of Na_V_1.5 voltage dependence.^*17-20*^ We tested each compound (10^−9^ to 10^−4^ M) for its ability to inhibit peak voltage-gated Na^+^ currents (Figure 1A-D). Of the agents tested, Triton X-100 (log*P*_OW_ 4.6, IC_50_ 5.3 µM, Figure 1C-D), was the most potent and capsaicin (log*P*_OW_ 3.04, IC_50_ 60.2 µM, Figure 1C-D, Table 1) the least. The antiarrythmic amiodarone (log*P*_OW_ 7.2, IC_50_ 8.4 µM, Figure 1C-D) and β-blocker propranolol (log*P*_OW_ 3.48, IC_50_ 7.6 µM, Figure 1C-D) also inhibited Na_V_1.5 voltage-gating. Propranolol, which had a partition coefficient (log*P*_OW_) similar to capsaicin, was an 8-fold more potent Na_V_1.5 inhibitor than the latter, indicating that log*P*_OW_ is not a good predictor of the drug’s effect on Na_V_1.5 voltage-gated function. Because our goal was to test mechanosensitivity while minimizing Na_V_1.5 voltage-dependent gating inhibition, we chose capsaicin (20 µM) for further investigation, as this dose inhibited voltage-dependent Na^+^ current by ≤25% without mechano stimulus (Figure 1B-D).

**Table 1.** Partition coefficients and IC_50_ values for amphipathic agents. Partition coefficients denoted log*P*_OW_ for amiodarone, capsaicin, propranolol and Triton-X100 were previously reported^*17-20*^. IC_50_, concentration at which an amphipathic agent inhibited half of the maximum peak whole cell Na^+^ current from HEK293 cells transfected with Na_V_1.5.

|  | Partition coefficient (log*P*<sub>OW</sub>) | IC<sub>50</sub> (μM) | Slope |
| --- | --- | --- | --- |
| Amiodarone | 7.2 | 8.4 | 0.50 |
| Capsaicin | 3.04 | 60 | 0.64 |
| Propranolol | 3.48 | 7.6 | 1.01 |
| Triton X-100 | 4.6 | 5.3 | 1.00 |

**Figure 1.**
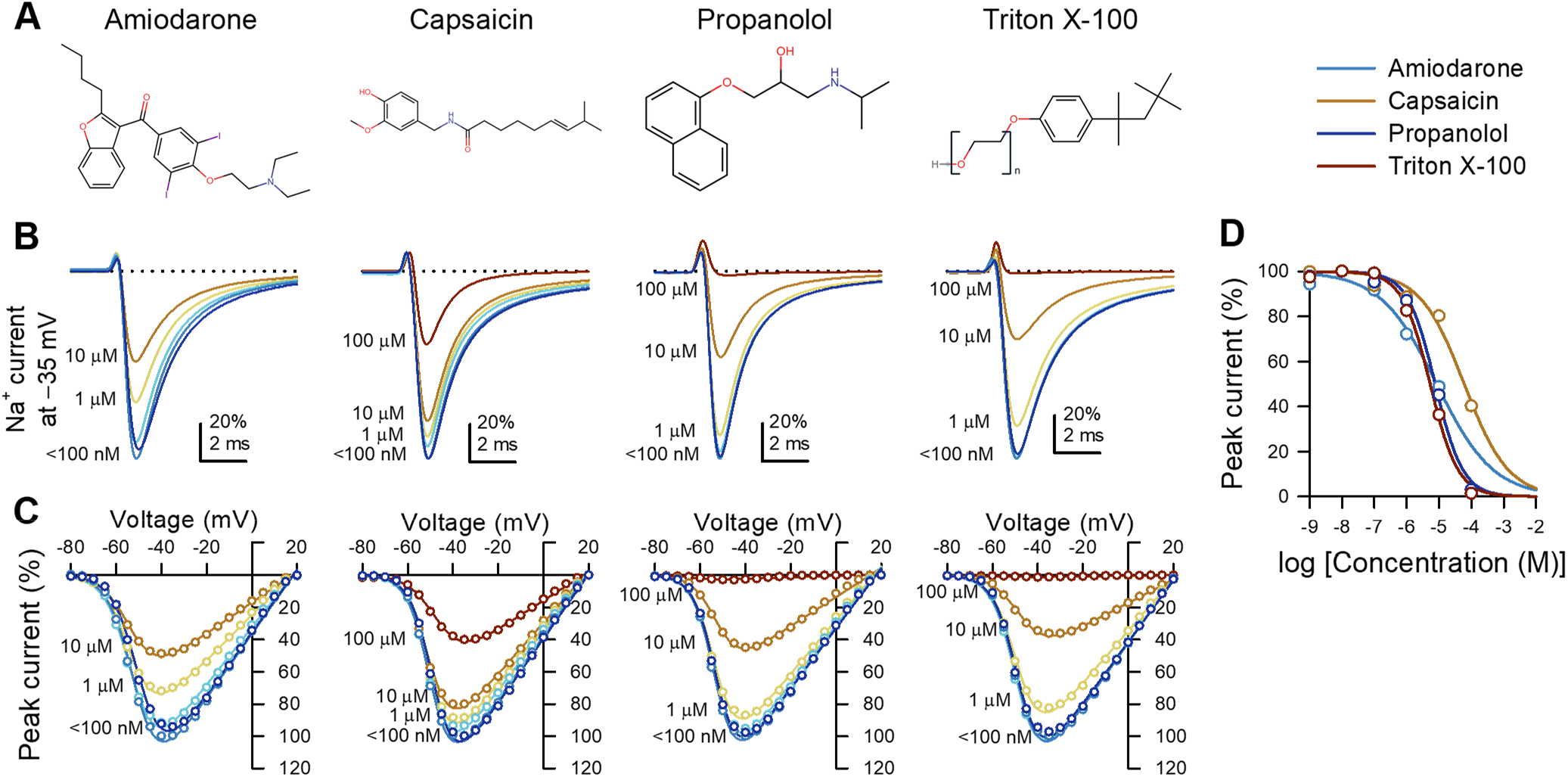
Amphipathic compounds inhibit voltage-gated Na^+^ currents from Na_V_1.5 channels expressed in HEK293 cells. *A*, Molecular structures of the amphipaths (from *left* to *right*): amiodarone, capsaicin, propranolol, and Triton X-100 .*B-C*, Representative Na^+^ currents elicited by a step from −120 to the −35-mV test voltage (*B*), and peak Na^+^ current-voltage plots across all test voltages (*C*) with 10^−9^ to 10^−4^ M (*blue-red spectrum*) of membrane-permeable amphipathic compounds in the extracellular solution. *D*, Dose-response curves for maximum peak Na^+^ current of Na_V_1.5 *vs*. amphipathic concentration; IC_50_ values: amiodarone, 8.4 µM; capsaicin, 60.2 µM; propranolol, 7.6 µM; Triton X-100, 5.3 µM.

### Capsaicin inhibits increases in peak current and acceleration with mechanical stimuli

To test the effect of capsaicin on Na_V_1.5 mechanosensitivity, we used two complementary approaches for mechanical stimulation: (1) cell-attached macroscopic patches with suction and (2) whole-cell configuration with fluid shear stress (Tables 2-3, Figure 2). These complementary techniques allow us to highlight different aspects of the channels’ mechanosensitivity due to both techniques’ intrinsic strengths and weaknesses.^*11, 21-23*^ The effect of pressure was tested in a pair-wise fashion^*1, 11, 14*^, with pressure at 0 or −30 mmHg applied at each voltage step (Figure 2A,C,E-H). Whole-cell current responses to shear was tested by perfusion at 0 or 10 mL/min (Figure 2B,D,E-H). We then reassessed function in both configurations in the presence of 20 µM capsaicin (Tables 2-3, Figure 2A-H). Suction increased normalized peak currents (I_MAX_) by 16.6±2.4% (*P*<0.05; n = 24; Figure 2A,C,E), and shear increased the peak current (I_PEAK_) by 16.0±3.1% in whole cells (0.26±0.10 nS increase in conductance; *P*<0.05, 0 to 10 mL/min; n = 12; Figure 2B,C,E). Capsaicin decreased I_PEAK_ by 22.1±3.9% (*P*<0.05, 0 to 20 µM capsaicin), and both pressure (+4.8±3.0%) and shear sensitivity (+3.1±3.8%, +0.08±0.05 nS) were lost (n = 12-14, *P*>0.05 to drug with no force).

**Table 2.** Effect of capsaicin on pressure-induced Na_V_1.5 mechanosensitivity in cell-attached patches. Effects of pressure (0 or −30 mmHg) on parameters of macroscopic Na^+^ currents without (0 µM) or with capsaicin (20 µM): maximum peak Na^+^ currents normalized to controls at 0 mmHg (I_MAX_), voltage dependence of activation (V_1/2A_) or inactivation (V_1/2I_), time constant of activation (τ_A_), time of inactivation recovery (t_1/2R_), slope of inactivation recovery (slope), maximum use-dependent inhibition (block), frequency of use-dependent inhibition (*f*_1/2_). n = 8-24 cells, \**P*<0.05, 0 to −30 mmHg or †*P*<0.05, 0 to 20 µM capsaicin by a 2-way ANOVA with Tukey post-test.

|  | 0 μM |  |  |  | 20 μM |  |  |  |
| --- | --- | --- | --- | --- | --- | --- | --- | --- |
|  | 0 mmHg | -30 mmHg | Change | n | 0 mmHg | -30 mmHg | Change | n |
| I<sub>MAX</sub> (%) | 100.0±0.0 | 116.6±2.4* | 16.6±2.4 | 24 | 100.0±0.0 | 104.8±3.0† | 4.8±3.0 | 14 |
| V<sub>1/2A</sub> (mV) | -41.9±3.0 | -46.4±3.0* | -4.5±0.6 | 24 | -57.5±2.8† | -60.0±3.1*† | -2.4±0.6 | 14 |
| V<sub>1/2I</sub> (mV) | -54.1±3.1 | -60.1±3.3* | -6.0±0.9 | 24 | -70.5±3.3† | -71.9±3.5† | -1.4±1.2 | 14 |
| τ<sub>A</sub> (ms) | 0.32±0.04 | 0.26±0.04* | -20.0±5.3% | 24 | 0.23±0.04 | 0.21±0.04 | -11.0±5.4% | 14 |
| t<sub>1/2R</sub> (ms) | 13.2±2.5 | 22.1±4.9* | 8.9±3.9 | 11 | 29.5±6.4† | 48.7±9.4*† | 19.2±8.6 | 11 |
| Slope (ms<sup>-1</sup>) | 1.3±0.1 | 1.2±0.1 | -0.10±0.17 | 11 | 1.9±0.3 | 2.3±0.8 | 0.41±0.87 | 11 |
| Block (%) | 80.9±7.9 | 77.8±8.6 | 3.2±9.4 | 11 | 93.6±4.5 | 85.5±8.5 | 4.7±3.8 | 8 |
| f<sub>1/2</sub> (Hz) | 22.3±2.6 | 20.0±2.1 | -2.4±3.3 | 11 | 19.1±2.5 | 14.8±1.4* | -4.2±1.4 | 8 |

**Figure 2.**
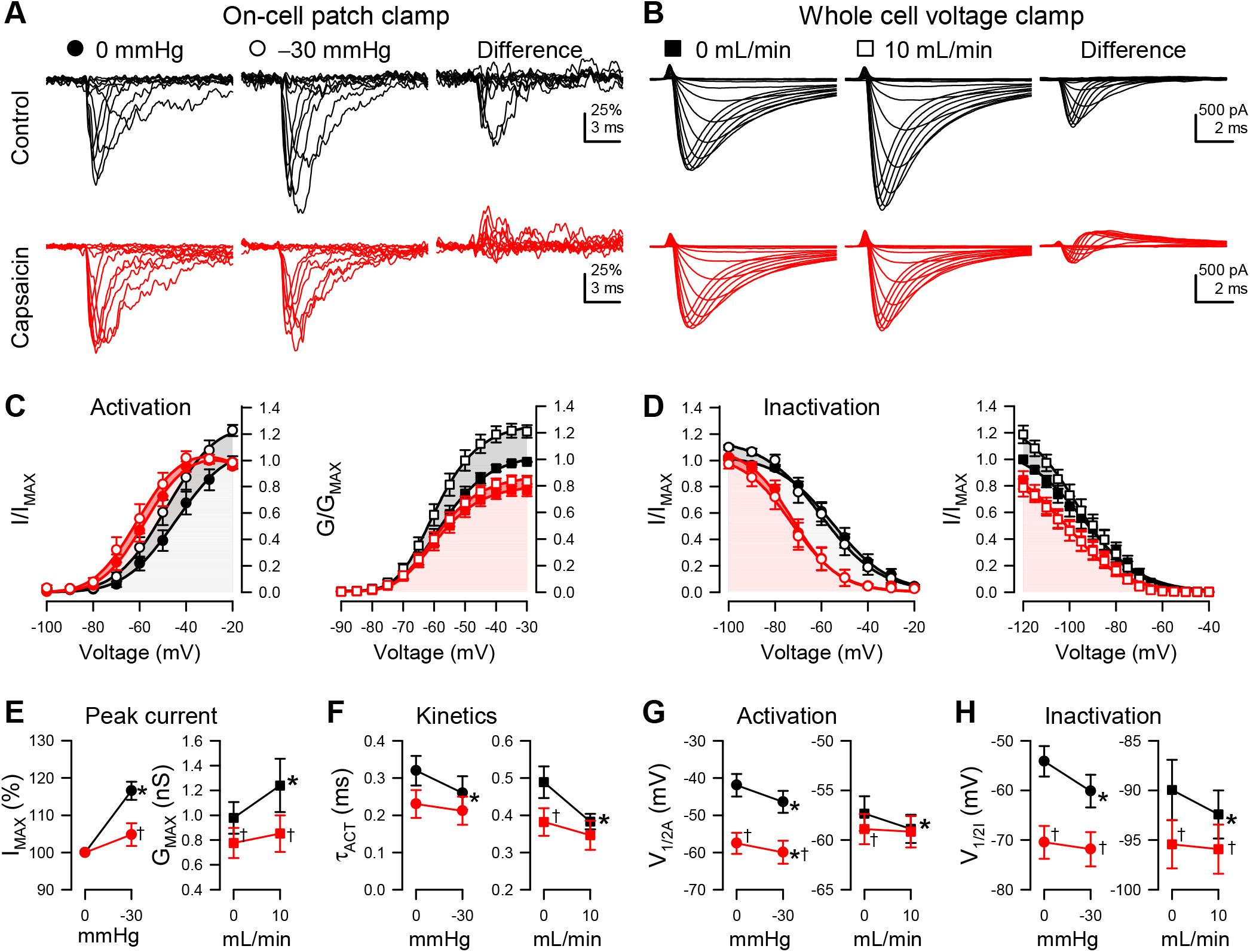
Capsaicin inhibits pressure- and shear-sensitivity of Na_V_1.5. *A*, Representative Na_V_1.5 currents elicited by voltage ladders ranging −100 to 0 mV in a cell-attached patch (*A*) or −120 mV to −30 mV in a whole cell (*B*), recorded at rest (*filled symbols*) or with force (*empty symbols*), in the presence of 0 µM (*black*) or 20 µM capsaicin (*red*). Difference currents were constructed by subtracting the control Na^+^ currents from the pressure-(*A*) or shear-stimulated (*B*) currents. *C-D*, Steady-state activation (*C*) and inactivation (*D*) curves of Na^+^ currents in cell-attached patches (*left*) or whole cells (*right*), recorded at rest (*filled symbols*) or with force (*empty symbols*), in the presence of 0 µM (*black*) or 20 µM capsaicin (*red*). *E-H*, Maximum peak Na^+^ current (*E*), time constant of activation (*F*), and voltage dependence of activation (*G*, V_1/2A_) or inactivation (*H*, V_1/2I_), recorded with 0 or −30 mmHg pressure in the patch (*left*) and 0 or 10 mL/min shear stress in whole cells (*right*) in the presence of 0 µM (*black*) or 20 µM capsaicin (*red*). n = 12-24 cells, \**P*<0.05 comparing 0 to −30 mmHg or 0 to 10 mL/min, †*P*<0.05 comparing 0 to 20 µM capsaicin by a 2-way ANOVA with Tukey post-test.

In the absence of drug, pressure and shear accelerated Na^+^ current activation, decreasing the activation constant (τ_ACT_) by 20.0±5.3% or 20.4±3.3%, respectively (n = 12-14, *P*<0.05 to no force controls; Figure 2F). Capsaicin accelerated Na_V_1.5 activation by 20.3±6.9% at rest (n = 12-14, *P*<0.05, 0 to 20 µM capsaicin) in whole cells but not in patches, and capsaicin inhibited the acceleration of activation induced by pressure and shear, as τ_ACT_ did not accelerate with pressure or shear (−11.0±5.4% or −1.3±7.0%, respectively; n = 12-14, *P*>0.05 to drug with no force, Figure 2F). Our results thus show that capsaicin inhibits the mechanosensitivity of peak current and kinetics of Na_V_1.5 in both experimental configurations.

### Capsaicin inhibits mechanically induced hyperpolarizing shifts in the voltage dependence of activation and availability

Pressure^*1, 13, 14, 24*^ and shear^*3, 11, 12*^ produce hyperpolarizing shifts in the voltage dependence of Na_V_1.5 activation and inactivation. Membrane-permeable amphipathic drugs like lidocaine and ranolazine reduce these mechanosensitive shifts in voltage dependence.^*13, 14*^ Therefore, we explored whether capsaicin could reduce the pressure- or shear-induced shifts in voltage dependence. Like in our previous work without drug^*13, 14*^, suction (−30 mmHg) produced a leftward shift of −4.5±0.6 mV, and shear stress induced a smaller but significant shift of −1.5±0.6 mV in the voltage dependence of activation (V_1/2A_) (*P*<0.05 to no force) (Table 2, Table 3, Figure 2C, G). Without force, capsaicin produced a hyperpolarized shift in V_1/2A_ (−1.6±0.4 mV; *P*<0.05, 0 to 20 µM capsaicin) in whole cells. With force, capsaicin inhibited the shear-induced shift in V_1/2A_ (−0.3±0.1 mV, *P*>0.05 to drug with no shear) but not the pressure-induced shift (−2.4±0.6 mV, *P*<0.05 to drug with no pressure). Similar to shear-induced shifts in V_1/2A_, pressure or shear shifted the voltage dependence of inactivation or availability (V_1/2I_) in the absence of capsaicin (−6.0±0.9 mV with pressure or −2.5±0.9 mV with shear, *P*<0.05 to no force) (Tables 2-3, Figure 2D,H). Without force, capsaicin produced a hyperpolarizing shift in whole-cell V_1/2I_ by −5.1±0.7 mV, as previously observed^*25*^. In the presence of capsaicin, however, neither pressure nor shear had a significant effect on V_1/2I_ (−1.4±1.2 or −0.5±0.6 mV change, respectively, *P*>0.05 to drug with no force), suggesting loss of the MS of Na_V_1.5 inactivation. Overall, our results show that capsaicin inhibited the mechanosensitive shifts in Na_V_1.5 gating.

**Table 3.** Effect of capsaicin on shear-induced Na_V_1.5 mechanosensitivity whole cells. Effects of shear stress (0 or 10 mL/min) on parameters of whole cell Na^+^ currents without (0 µM) or with capsaicin (20 µM): maximum peak conductance (G_MAX_), maximum peak current density (I_PEAK_), voltage dependence of activation (V_1/2A_) or inactivation (V_1/2I_), time constants of activation (τ_A_) and fast (τ_F_) or slow inactivation (τ_S_), time of inactivation recovery (t_1/2R_), slope of inactivation recovery (slope), maximum use-dependent inhibition (block), frequency of use-dependent inhibition (*f*_1/2_). n = 8-18 cells, \**P*<0.05, 0 to 10 mL/min or †*P*<0.05, 0 to 20 µM capsaicin by a 2-way ANOVA with Tukey post-test.

|  | 0 μM |  |  |  | 20 μM |  |  |  |
| --- | --- | --- | --- | --- | --- | --- | --- | --- |
|  | 0 mL/min | 10 mL/min | Change | n | 0 mL/min | 10 mL/min | Change | n |
| **G<sub>MAX</sub> (nS)** | 0.98±0.13 | 1.24±0.22* | 0.26±0.10 | 12 | 0.78±0.12<sup>†</sup> | 0.85±0.15<sup>†</sup> | 0.08±0.05 | 12 |
| **I<sub>PEAK</sub> (pA/pF)** | -66.8±12.3 | -86.6±17.7* | 16.0±3.1% | 12 | -56.7±58.5<sup>†</sup> | -58.5±11.6<sup>†</sup> | 3.1±3.8%<sup>†</sup> | 12 |
| **V<sub>1/2A</sub> (mV)** | -57.3±1.7 | -58.9±1.4* | -1.5±0.6 | 12 | -58.9±1.5<sup>†</sup> | -59.2±1.6 | -0.3±0.1 | 12 |
| **V<sub>1/2I</sub> (mV)** | -90.3±3.3 | -92.7±2.6* | -2.5±0.9 | 12 | -95.4±2.6<sup>†</sup> | -95.8±2.7<sup>†</sup> | -0.5±0.6 | 12 |
| **τ<sub>A</sub> (ms)** | 0.5±0.05 | 0.39±0.02* | -20.4±3.3% | 12 | 0.39±0.04<sup>†</sup> | 0.38±0.04 | -1.3±7.0%<sup>†</sup> | 12 |
| **τ<sub>F</sub> (ms)** | 0.82±0.08 | 0.57±0.04* | -27.5±4.0% | 12 | 0.80±0.08 | 0.56±0.05* | -25.1±5.9% | 12 |
| **τ<sub>S</sub> (ms)** | 4.7±0.3 | 3.5±0.2* | -24.2±4.4% | 12 | 3.8±0.3<sup>†</sup> | 2.9±0.2*<sup>†</sup> | -20.5±4.8% | 12 |
| **t<sub>1/2R</sub> (ms)** | 18.8±1.7 | 16.6±1.8* | -2.2±0.6 | 8 | 38.0±4.5<sup>†</sup> | 28.4±4.2*<sup>†</sup> | -9.5±0.9<sup>†</sup> | 8 |
| **slope (ms<sup>-1</sup>)** | -1.26±0.04 | -1.28±0.05 | -0.02±0.02 | 8 | -1.43±0.02<sup>†</sup> | -1.42±0.06 | 0.01±0.04 | 8 |
| **Block (%)** | 64.6±2.4 | 64.7±2.6 | 0.2±1.7 | 18 | 89.0±1.4<sup>†</sup> | 83.9±2.0*<sup>†</sup> | -5.1±0.9<sup>†</sup> | 10 |
| **f<sub>1/2</sub> (Hz)** | 26.1±2.0 | 27.3±2.3 | 1.2±0.9 | 18 | 18.9±1.0<sup>†</sup> | 21.0±1.7 | 2.1±1.4 | 10 |

### Effects of capsaicin and mechanical stimuli on recovery from inactivation

Both capsaicin and pressure delay recovery of Na_V_1.5 from fast inactivation.^***1, 25***^ Therefore, we tested whether the presence of capsaicin affected the recovery from fast inactivation (1 to 1000 ms) in the absence or presence of mechanical stimuli (Tables 2-3, Figure 3A-B). Without force or drug, Na^+^ currents recovered within ∼100 ms in either configuration (Figure 3C); the half-time of Na_V_1.5 inactivation recovery (t_1/2R_) at rest was 13.2±2.5 ms in the patch and 18.8±1.7 ms in whole-cell (Tables 2-3, Figure 3C-F). In addition, unlike the consistent responses to force regardless of stimulus or configuration described above, here we observed consistent differences between the two approaches. Shear accelerated Na_V_1.5 t_1/2R_ by 2.2±0.6 ms (*P*<0.05, 0 to 10 mL/min), whereas pressure delayed the t_1/2R_ (+8.9±3.9 ms, *P*<0.05, 0 to −30 mmHg) (Tables 2-3, Figure 3C-F). In whole cells, without force, capsaicin delayed the recovery from inactivation; t_1/2R_ increased from 18.8±1.7 to 38.0±4.5 ms (*P*<0.05, 0 to 20 µM capsaicin). With capsaicin present, pressure increased t_1/2R_ by 19.2±8.2 ms (*P*<0.05 to drug with no pressure), whereas shear reduced t_1/2R_ in whole cells by 9.5±0.9 ms (*P*<0.05 to drug with no shear). These data show that shear stress on whole cells and pressure on patches had opposite effects (shear accelerating, suction slowing). In both approaches, the recovery from inactivation was delayed by capsaicin, and capsaicin further delayed recovery in patches but accelerated recovery in whole cells.

**Figure 3.**
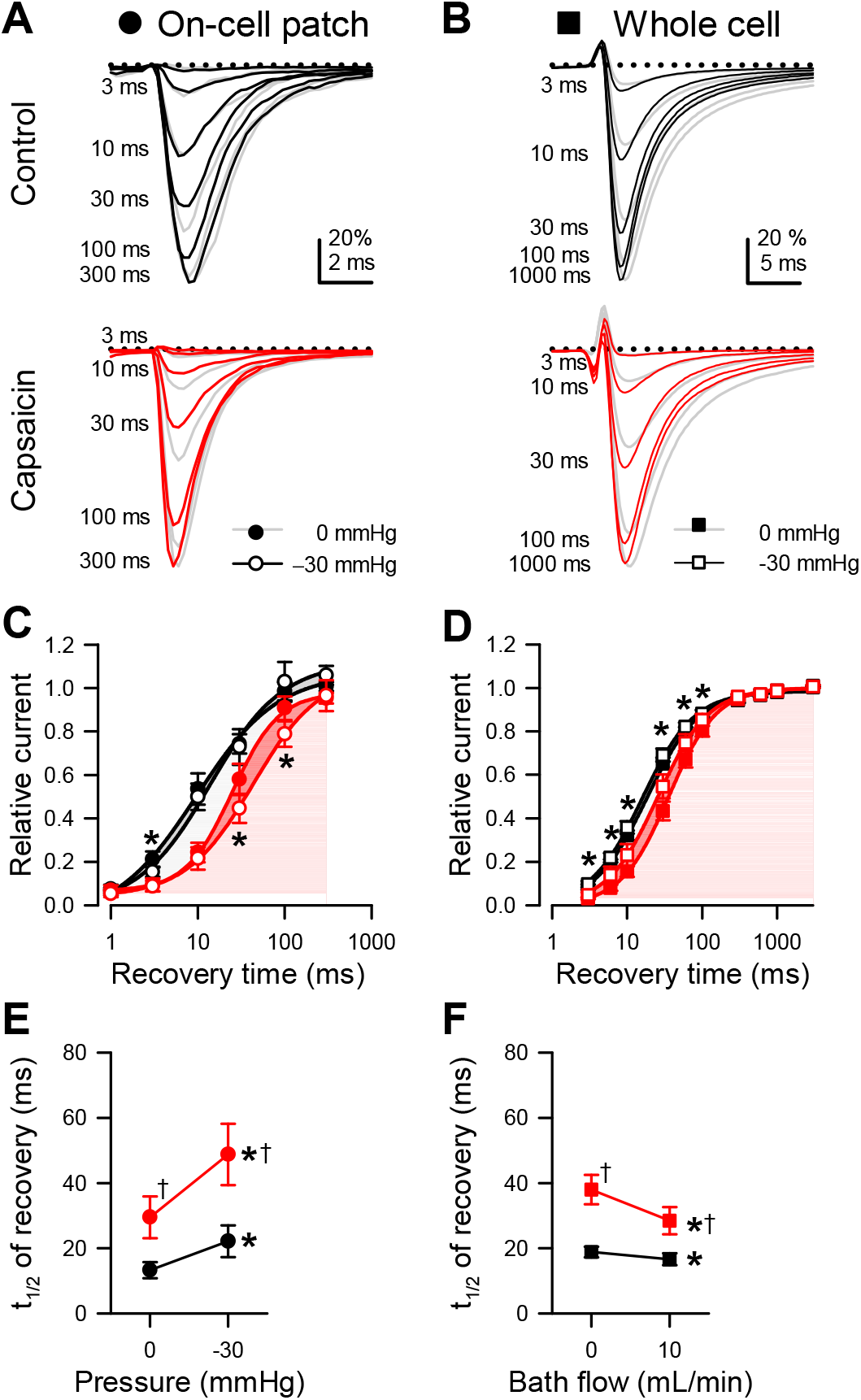
Effects of capsaicin on mechanosensitivity of Na_V_1.5 inactivation recovery time. *A-B*, Representative Na_V_1.5 currents at −20 mV in a cell-attached patch (*A*, **•**) or −30 mV in a whole cell (*B*, **■**), elicited after recovering from the control pulse for 3-300 ms at −120 mV (*A*) or 3-1000 ms at −130 mV (*B*). Na^+^ currents were recorded at rest (*grey*) or with force (*dark traces*: *A*, −30 mmHg pressure; *B*, 10 mL/min shear stress) in the presence of 0 µM (*top*) or 20 µM capsaicin (*bottom*). *C-D*, Normalized peak Na^+^ current versus recovery time in the presence of 0 µM (*black*) or 20 µM capsaicin (*red*), at 0 (**•**) or −30 mmHg pressure (○) in the patch (*C*) or at 0 (**■**) or 10 mL/min (□) shear stress in whole cells (*D*). *E-F*, Inactivation recovery times (t_1/2_) versus 0 or −30 mmHg pressure in the patch (*E*) and 0 or 10 mL/min shear stress in whole cells (*F*) with 0 µM (*black*) or 20 µM capsaicin (*red*). n = 8-11 cells, \**P*<0.05 comparing 0 to −30 mmHg or 0 to 10 mL/min, †*P*<0.05 comparing 0 to 20 µM capsaicin by a 2-way ANOVA with Tukey post-test.

### Effects of capsaicin and mechanical stimuli on use-dependent inactivation

Capsaicin can stabilize the inactivated state of Na_V_1.5 through use-dependent inhibition.^*25*^ The effects of pressure on use-dependent Na_V_1.5 function have not been fully explored, and we tested whether force can alter use-dependent inactivation of Na_V_1.5 in the absence or presence of capsaicin (Figure 4A-F). To measure the use-dependent inhibition of Na_V_1.5 expressed in HEK cells, Na^+^ currents elicited by steps to 0 or −30 mV in patches or whole cells were sampled at 3-33 Hz or 0.3-50 Hz, resepectively. Without force or drug, the maximum use-dependent inhibition of Na_V_1.5 was 80.9±7.9% with a half-frequency (*f*_1/2_) of 22.3±2.6 Hz in patches (Table 2, Figure 4C,E-F), and 64.6±2.4% with a *f*_1/2_ of 26.1±2.0 Hz in whole-cells (Table 3, Figure 4D-F). The use dependence did not change with either pressure or shear in the absence of capsaicin (*P*>0.05 to no force, Tables 2-3, Figure 4C-F). In the absence of shear, capsaicin increased the maximum use-dependent inhibition of Na_V_1.5 to 89.0±1.4% and decreased *f*_1/2_ to 18.9±1.0 Hz (*P*<0.05, 0 to 20 µM capsaicin). In the presence of capsaicin, shear produced a modest decrease in the maximum use-dependent inhibition (5.1±0.9%; *P*<0.05 to drug with no shear), and *f*_1/2_ was unaffected, suggesting that shear partially reverses the use-dependent inhibition of Na_V_1.5 promoted by capsaicin. In patches, capsaicin affected neither the use-dependent inhibition nor *f*_1/2_ at rest (*P*>0.05, 0 to 20 µM capsaicin) but increased the pressure-sensitivity (*f*_1/2_ decreased by 2.4±3.3 Hz; *P*<0.05 to drug with no pressure), suggesting that pressure is synergestic with capsaicin to decrease the frequency at which Na_V_1.5 undergoes use dependent inhibition. Together, our results suggest that though capsaicin enhances use-dependent inhibition, its effect on force-dependent changes to Na_V_1.5 use dependence may be specific to the type of force applied.

**Figure 4.**
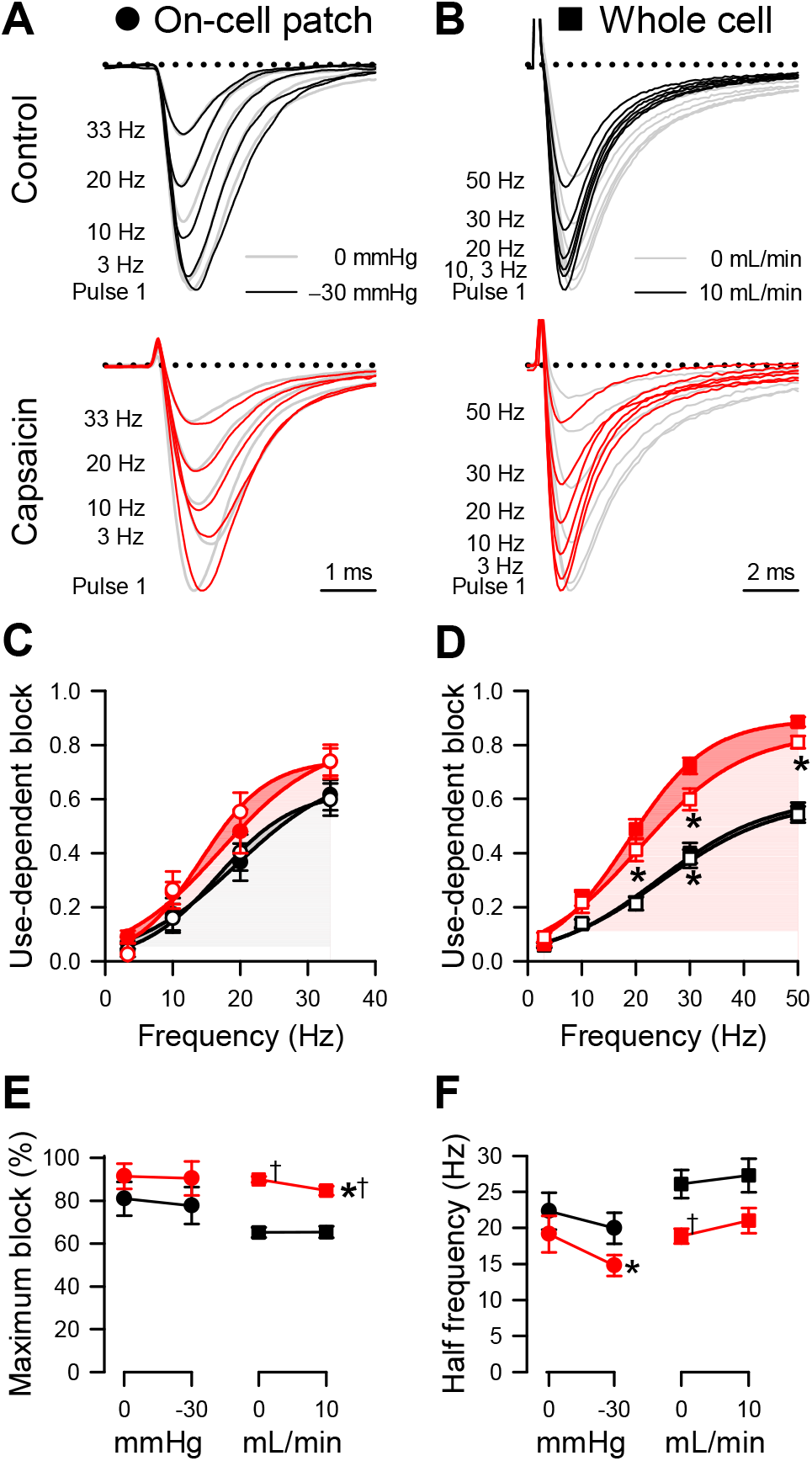
Effects of capsaicin on mechanosensitivity of Na_V_1.5 use-dependent inactivation. *A-B*, Representative Na_V_1.5 currents at the 20^th^ pulse to −20 mV in a cell-attached patch (*A*, **•**) or to −40 mV in a whole cell (*B*, **■**), elicited at interpulse frequencies 3-33 Hz (*A*) or 3-50 Hz (*B*). Na^+^ currents were recorded at rest (*grey*) or with force (*dark traces*: *A*, −30 mmHg pressure; *B*, 10 mL/min shear stress) in the presence of 0 µM (*top*) or 20 µM capsaicin (*bottom*). *C-D*, Use-dependent inhibition of peak Na^+^ current versus interpulse frequency in the presence of 0 µM (*black*) or 20 µM capsaicin (*red*), at 0 (**•**) or −30 mmHg pressure (○) in the patch (*C*) or at 0 (**■**) or 10 mL/min (□) shear stress in whole cells (*D*). *E-F*, Maximum use-dependent inhibition (*E*) or frequency of use-dependent inhibition (*F*) versus pressure in the patch (*left*) and shear stress in whole cells (*right*) with 0 µM (*black*) or 20 µM capsaicin (*red*). n = 8-18 cells, \**P*<0.05 comparing 0 to −30 mmHg or 0 to 10 mL/min, †*P*<0.05 comparing 0 to 20 µM capsaicin by a 2-way ANOVA with Tukey post-test.

## DISCUSSION

Our study aimed to test the impact of a membrane-permeable amphipathic agent on Na_V_1.5 mechanosensitivity. We selected compounds with high partition coefficients and tested the inhibition of Na_V_1.5 voltage-gating. Capsaicin inhibited Na_V_1.5 mechanosensitivity comparable to the amphipaths lidocaine^*13, 14*^ and ranolazine^*14*^. Capsaicin consistently inhibited the effects of pressure or shear stress on Na_V_1.5 in membrane patches or whole cells, respectively, by (1) diminishing the mechanosensitive increases in Na^+^ current, (2) shifts in steady-state voltage dependence, and (3) acceleration of Na_V_1.5 gating kinetics.

Quantifying mechanosensitive changes in ion channel function can be challenging, and few studies have explored Na_V_1.5 mechanosensitivity using whole-cell and patch modes in parallel.^*1,11, 13*^ To our knowledge, this is the first study to compare the effects of pressure and shear on the mechanosensitive operation of Na_V_1.5 in the absence nor presence of drug. Most of Na_V_1.5’s mechanosensitive responses and capsaicin’s effects on Na_V_1.5 MS were similar in the two testing modes. Both produced an increase in peak Na^+^ current, a shift in V_1/2A_, a shift in V_1/2I_, and an acceleration in τ_A_. Capsaicin, using either approach, inhibited MS effects on the above biophysical parameters. This is important because mechanical strain leads to faster and greater Na^+^ influx, which increases Na_V_ channel availability and further depolarizes the membrane (closer to the threshold to fire action potentials or elicit autonomous membrane depolarizations). Capsaicin would reduce Na^+^ influx, slow membrane depolarization (or hyperpolarize the membrane) thereby reducing the effect of mechanical forces on Na_V_ channel availability.

Surprisingly, we found opposite responses in the pressure- and shear-sensitivity of Na_V_1.5 inactivation recovery and use-dependent inactivation. Pressure increased the inactivation recovery time, whereas shear stress decreased it. Addition of capsaicin to patches resulted in a further delay of Na_V_1.5 inactivation recovery in patches under pressure, in contrast to whole cells, where shear stress accelerated inactivation recovery whether or not capsaicin was present. Similarly, capsaicin and pressure, when applied together, decreased the frequency of use-dependent inactivation in patches, while in whole cells capsaicin alone lowered use-dependence frequency, a process unaffected by shear stress. Pressure has previously been shown to prolong Na_V_1.5 inactivation recovery time in patches^1^, but the effect of shear stress on inactivation recovery or use-dependent inhibition of Na_V_1.5 was previously unknown. The opposite responses in use dependence and recovery using the two approaches are independent of capsaicin, and therefore suggest that these mechanical stimuli may act through different mechanisms.^*21, 26*^ Conceivably, the effect of pressure or shear stress on the membrane or cytoskeleton could be different. The similar increases in peak current, as a result of pressure or shear, reflect a greater probability of Na_V_ channels in the open state, whereas the divergent effects of pressure and shear on use-dependence and inactivation recovery represent different pathways by which Na_V_ channels are entering or leaving the inactivated state. Shear can lead to uniaxial elastic tension along the membrane, yielding asymmetrical sliding of lipid membrane leaflets^29,30^, and the effects of lipid bilayer thinning affect some functional modes, such as inactivation, more than others^*27, 28*^. Meanwhile, macroscopic patch suction can create unequal transmembrane surface tension^*2, 21, 29*^, with the tension being greatest at the top of the dome. Therefore, while both stimuli may be used to study mechanosensitivity of Na_V_1.5 and other mechanosensitive ion channels, functional consequences may be dependent on channel’s functional modes under study.

Capsaicin can inhibit Na_V_1.5 at rest by promoting Na_V_ channel inactivation but shows greater potency for inhibiting the inactivation-removed (IR) Na_V_ channel triple mutant, WCW, on the pore lumen DIS6 segment, in the analogous position and across from F1760 in the local anesthetic binding site on the DIVS6 segment.^*25, 30*^ Interestingly, the IR sequence in Na_V_1.5 (L407W-L409C-A410W) would be 9 amino acids downstream from and functionally similar to IR mutant T220A in bacterial homolog NaChBac, which has greater shear-sensitivity than its wild-type. Whether inhibition of mechanosensitivity by capsaicin requires the LXLA sequence in Na_V_1.5 remains to be seen, yet finding a shared interaction motif suggests that the LA binding region might sensitize Na_V_1.5 and other mechanosensitive voltage-gated channels^*31*^ or mechano-gated channels^*32*^ to amphipathic MS inhibition. Capsaicin had divergent effects on pressure- and shear-sensitivity of Na_V_1.5 use-dependent inhibition or inactivation recovery, suggesting that specific types of force may differentially modulate Na_V_1.5 mechanosensitivity.

Another amphiphilic Na_V_1.5 modulator, ranolazine, inhibits the increase in peak Na^+^ current and the hyperpolarization of voltage dependence of activation induced by pressure or shear stress comparable to capsaicin^*14*^. Ranolazine is an anti-ischemic agent that may cause constipation as a common side effect ^*33*^; muscle contractility in human colon smooth muscle cells was lost when ranolazine inhibited Na_V_1.5 peak current and mechanosensitivity.^*34*^ The effects of capsaicin, lidocaine^*13*^, and ranolazine^*14*^ as mechanosensitivity inhibitors demonstrate that membrane-permeable amphipathic agents may be candidates for modulating Na_V_1.5 mechanosensitivity and targeting dysfunction in mechanosensitive channelopathies. Amphipathic drugs are widely used in the clinical practice to target ion channels, but are rarely used for mechano-modulation.^12,13^ Channelopathies involving mechanosensitive dysfunction are an emerging area of study.^*6, 7, 9-11, 35*^ Ion channelopathies in voltage-sensitive mechano-gated Piezo channels ^*36-38*^ and channelopathies affecting Na_V_1.5 mechanosensitivity have been identified, but both lack treatment options. Hence, drugs that can target and modulate mechanosensitivity carry promise for treating disease.

Capsaicin inhibits its canonical target TRPV1 in sensory neurons to improve GI dysfunction in IBS-D patients^*39*^; however, TRPV1 is not pressure-sensitive up to −90 mmHg^*40*^ and does not have high expression in HEK cells^*29, 41*^. Capsaicin has shown promise in targeting pain in IBS. Interestingly, it also has effects on gut motility^*42-44*^, possibly through it’s function on Na_V_1.5 mechanosensivity. Therefore, there may be an exciting possibility of using capsaicin to affect sensory (TRPV1) and motility (SCN5A/Na_V_1.5) processes by different mechanisms in the GI tract.

## ACKNOWLEDGEMENTS

We thank Kristy Zodrow for administrative assistance. This work was supported by NIH grants DK052766 (GF), DK106456 (AB), and the National Center for Complementary and Integrative Health AT10875 (AB).

## REFERENCES

[1] Beyder, A., Rae, J. L., Bernard, C., Strege, P. R., Sachs, F., and Farrugia, G. (2010) Mechanosensitivity of Nav1.5, a voltage-sensitive sodium channel, J Physiol 588, 4969–4985.

[2] Sukharev, S., and Sachs, F. (2012) Molecular force transduction by ion channels: diversity and unifying principles, J Cell Sci 125, 3075–3083.

[3] Strege, P. R., Holm, A. N., Rich, A., Miller, S. M., Ou, Y., Sarr, M. G., and Farrugia, G. (2003) Cytoskeletal modulation of sodium current in human jejunal circular smooth muscle cells, Am J Physiol Cell Physiol 284, C60–66.

[4] Beyder, A., and Farrugia, G. (2016) Ion channelopathies in functional GI disorders, Am. J. Physiol. Gastrointest. Liver Physiol. 311, G581–G586.

[5] Kass, R. S. (2005) The channelopathies: novel insights into molecular and genetic mechanisms of human disease, The Journal of clinical investigation 115, 1986–1989.

[6] Marban, E. (2002) Cardiac channelopathies, Nature 415, 213–218.

[7] Beyder, A., and Farrugia, G. (2016) Ion channelopathies in functional GI disorders, Am J Physiol Gastrointest Liver Physiol 311, G581–G586.

[8] Banderali, U., Juranka, P. F., Clark, R. B., Giles, W. R., and Morris, C. E. (2010) Impaired stretch modulation in potentially lethal cardiac sodium channel mutants, Channels (Austin) 4, 12–21.

[9] Beyder, A., Mazzone, A., Strege, P. R., Tester, D. J., Saito, Y. A., Bernard, C. E., Enders, F. T., Ek, W. E., Schmidt, P. T., Dlugosz, A., Lindberg, G., Karling, P., Ohlsson, B., Gazouli, M., Nardone, G., Cuomo, R., Usai-Satta, P., Galeazzi, F., Neri, M., Portincasa, P., Bellini, M., Barbara, G., Camilleri, M., Locke, G. R., Talley, N. J., D’Amato, M., Ackerman, M. J., and Farrugia, G. (2014) Loss-of-function of the voltage-gated sodium channel NaV1.5 (channelopathies) in patients with irritable bowel syndrome, Gastroenterology 146, 1659–1668.

[10] Saito, Y. A., Strege, P. R., Tester, D. J., Locke, G. R., 3rd, Talley, N. J., Bernard, C. E., Rae, J. L., Makielski, J. C., Ackerman, M. J., and Farrugia, G. (2009) Sodium channel mutation in irritable bowel syndrome: evidence for an ion channelopathy, Am. J. Physiol. Gastrointest. Liver Physiol. 296, G211–218.

[11] Strege, P. R., Mercado-Perez, A., Mazzone, A., Saito, Y. A., Bernard, C. E., Farrugia, G., and Beyder, A. (2019) SCN5A mutation G615E results in NaV1.5 voltage-gated sodium channels with normal voltage-dependent function yet loss of mechanosensitivity, Channels (Austin) 13, 287–298.

[12] Strege, P. R., Mazzone, A., Bernard, C. E., Neshatian, L., Gibbons, S. J., Saito, Y. A., Tester, D. J., Calvert, M. L., Mayer, E. A., Chang, L., Ackerman, M. J., Beyder, A., and Farrugia, G. (2017) Irritable bowel syndrome (IBS) patients have SCN5A channelopathies that lead to decreased NaV1.5 current and mechanosensitivity, Am J Physiol Gastrointest Liver Physiol 314, G494–G503.

[13] Beyder, A., Strege, P. R., Bernard, C., and Farrugia, G. (2012) Membrane permeable local anesthetics modulate Na(V)1.5 mechanosensitivity, Channels (Austin) 6, 308–316.

[14] Beyder, A., Strege, P. R., Reyes, S., Bernard, C. E., Terzic, A., Makielski, J., Ackerman, M. J., and Farrugia, G. (2012) Ranolazine decreases mechanosensitivity of the voltage-gated sodium ion channel Na(v)1.5: a novel mechanism of drug action, Circulation 125, 2698–2706.

[15] Kraichely, R. E., Strege, P. R., Sarr, M. G., Kendrick, M. L., and Farrugia, G. (2009) Lysophosphatidyl choline modulates mechanosensitive L-type Ca2+ current in circular smooth muscle cells from human jejunum, Am J Physiol Gastrointest Liver Physiol 296, G833–839.

[16] Makielski, J. C., Ye, B., Valdivia, C. R., Pagel, M. D., Pu, J., Tester, D. J., and Ackerman, M. J. (2003) A ubiquitous splice variant and a common polymorphism affect heterologous expression of recombinant human SCN5A heart sodium channels, Circ. Res. 93, 821–828.

[17] Ho, Y. F., Chou, H. Y., Chu, J. S., and Lee, P. I. (2018) Comedication with interacting drugs predisposes amiodarone users in cardiac and surgical intensive care units to acute liver injury: A retrospective analysis, Medicine (Baltimore) 97, e12301.

[18] LaHann, T. R., DeKrey, L. J., and Tarr, B. D. (1989) Capsaicin analgesia: predictions based on physico-chemical properties, Proc West Pharmacol Soc 32, 201–204.

[19] Avdeef, A., Box, K. J., Comer, J. E., Hibbert, C., and Tam, K. Y. (1998) pH-metric logP 10. Determination of liposomal membrane-water partition coefficients of ionizable drugs, Pharm Res 15, 209–215.

[20] PubChem. (2019) PubChem Compound Summary for CID 5590, Octoxinol, NIH, National Library of Medicine.

[21] Suchyna, T. M., Markin, V. S., and Sachs, F. (2009) Biophysics and structure of the patch and the gigaseal, Biophys J 97, 738–747.

[22] Morris, C. E. (2011) Voltage-gated channel mechanosensitivity: fact or friction?, Front Physiol 2, 25.

[23] Sokabe, M., and Sachs, F. (1990) The structure and dynamics of patch-clamped membranes: a study using differential interference contrast light microscopy, J Cell Biol 111, 599–606.

[24] Morris, C. E., and Juranka, P. F. (2007) Nav channel mechanosensitivity: activation and inactivation accelerate reversibly with stretch, Biophys. J. 93, 822–833.

[25] Lundbaek, J. A., Birn, P., Tape, S. E., Toombes, G. E., Sogaard, R., Koeppe, R. E., 2nd, Gruner, S. M., Hansen, A. J., and Andersen, O. S. (2005) Capsaicin regulates voltage-dependent sodium channels by altering lipid bilayer elasticity, Mol Pharmacol 68, 680–689.

[26] Dimitrakopoulos, P. (2012) Analysis of the variation in the determination of the shear modulus of the erythrocyte membrane: Effects of the constitutive law and membrane modeling, Phys Rev E Stat Nonlin Soft Matter Phys 85, 041917.

[27] Lundbaek, J. A., Koeppe, R. E., 2nd, and Andersen, O. S. (2010) Amphiphile regulation of ion channel function by changes in the bilayer spring constant, Proc. Natl. Acad. Sci. U. S. A. 107, 15427–15430.

[28] Lundbaek, J. A., Collingwood, S. A., Ingolfsson, H. I., Kapoor, R., and Andersen, O. S. (2010) Lipid bilayer regulation of membrane protein function: gramicidin channels as molecular force probes, J R Soc Interface 7, 373–395.

[29] Bavi, N., Nakayama, Y., Bavi, O., Cox, C. D., Qin, Q. H., and Martinac, B. (2014) Biophysical implications of lipid bilayer rheometry for mechanosensitive channels, Proc Natl Acad Sci U S A 111, 13864–13869.

[30] Wang, S. Y., Mitchell, J., and Wang, G. K. (2007) Preferential block of inactivation-deficient Na+ currents by capsaicin reveals a non-TRPV1 receptor within the Na+ channel, Pain 127, 73–83.

[31] Lyford, G. L., Strege, P. R., Shepard, A., Ou, Y., Ermilov, L., Miller, S. M., Gibbons, S. J., Rae, J. L., Szurszewski, J. H., and Farrugia, G. (2002) alpha(1C) (Ca(V)1.2) L-type calcium channel mediates mechanosensitive calcium regulation, Am J Physiol Cell Physiol 283, C1001–1008.

[32] Joshi, V., Strege, P. R., Farrugia, G., and Beyder, A. (2021) Mechanotransduction in gastrointestinal smooth muscle cells: role of mechanosensitive ion channels, Am J Physiol Gastrointest Liver Physiol 320, G897–G906.

[33] Nash, D. T., and Nash, S. D. (2008) Ranolazine for chronic stable angina, Lancet 372, 1335–1341.

[34] Neshatian, L., Strege, P. R., Rhee, P. L., Kraichely, R. E., Mazzone, A., Bernard, C. E., Cima, R. R., Larson, D. W., Dozois, E. J., Kline, C. F., Mohler, P. J., Beyder, A., and Farrugia, G. (2015) Ranolazine inhibits voltage-gated mechanosensitive sodium channels in human colon circular smooth muscle cells, Am J Physiol Gastrointest Liver Physiol 309, G506–512.

[35] Locke, G. R., 3rd, Ackerman, M. J., Zinsmeister, A. R., Thapa, P., and Farrugia, G. (2006) Gastrointestinal symptoms in families of patients with an SCN5A-encoded cardiac channelopathy: evidence of an intestinal channelopathy, Am J Gastroenterol 101, 1299–1304.

[36] Alper, S. L. (2017) Genetic Diseases of PIEZO1 and PIEZO2 Dysfunction, Curr Top Membr 79, 97–134.

[37] Zarychanski, R., Schulz, V. P., Houston, B. L., Maksimova, Y., Houston, D. S., Smith, B., Rinehart, J., and Gallagher, P. G. (2012) Mutations in the mechanotransduction protein PIEZO1 are associated with hereditary xerocytosis, Blood 120, 1908–1915.

[38] Bae, C., Gnanasambandam, R., Nicolai, C., Sachs, F., and Gottlieb, P. A. (2013) Xerocytosis is caused by mutations that alter the kinetics of the mechanosensitive channel PIEZO1, Proc Natl Acad Sci U S A 110, E1162–1168.

[39] Gonlachanvit, S., Mahayosnond, A., and Kullavanijaya, P. (2009) Effects of chili on postprandial gastrointestinal symptoms in diarrhoea predominant irritable bowel syndrome: evidence for capsaicin-sensitive visceral nociception hypersensitivity, Neurogastroenterol Motil 21, 23–32.

[40] Nikolaev, Y. A., Cox, C. D., Ridone, P., Rohde, P. R., Cordero-Morales, J. F., Vasquez, V., Laver, D. R., and Martinac, B. (2019) Mammalian TRP ion channels are insensitive to membrane stretch, J Cell Sci 132.

[41] Mazzone, A., Gibbons, S. J., Eisenman, S. T., Strege, P. R., Zheng, T., D’Amato, M., Ordog, T., Fernandez-Zapico, M. E., and Farrugia, G. (2019) Direct repression of anoctamin 1 (ANO1) gene transcription by Gli proteins, FASEB J 33, 6632–6642.

[42] Agarwal, M. K., Bhatia, S. J., Desai, S. A., Bhure, U., and Melgiri, S. (2002) Effect of red chillies on small bowel and colonic transit and rectal sensitivity in men with irritable bowel syndrome, Indian J. Gastroenterol. 21, 179–182.

[43] Aniwan, S., and Gonlachanvit, S. (2014) Effects of Chili Treatment on Gastrointestinal and Rectal Sensation in Diarrhea-predominant Irritable Bowel Syndrome: A Randomized, Double-blinded, Crossover Study, J. Neurogastroenterol. Motil. 20, 400–406.

[44] Patcharatrakul, T., and Gonlachanvit, S. (2016) Chili Peppers, Curcumins, and Prebiotics in Gastrointestinal Health and Disease, Curr Gastroenterol Rep 18, 19.

